# Real-time monitoring and analysis of SARS-CoV-2 nanopore sequencing with minoTour

**DOI:** 10.1101/2021.09.13.459777

**Authors:** Rory Munro, Nadine Holmes, Christopher Moore, Matt Carlile, Alexander Payne, Roberto Santos, Matt Loose

## Abstract

**Motivation:** The ongoing SARS-CoV-2 pandemic has demonstrated the utility of real-time analysis of sequencing data, with a wide range of databases and resources for analysis now available. Here we show how the real-time nature of Oxford Nanopore Technologies sequencers can accelerate consensus generation, lineage and variant status assignment. We exploit the fact that multiplexed viral sequencing libraries quickly generate sufficient data for the majority of samples, with diminishing returns on remaining samples as the sequencing run progresses. We demonstrate methods to determine when a sequencing run has passed this point in order to reduce the time required and cost of sequencing.

**Results:** We extended MinoTour, our real-time analysis and monitoring platform for nanopore sequencers, to provide SARS-CoV2 analysis using ARTIC network pipelines. We additionally developed an algorithm to predict which samples will achieve sufficient coverage, automatically running the ARTIC medaka informatics pipeline once specific coverage thresholds have been reached on these samples. After testing on run data, we find significant run time savings are possible, enabling flow cells to be used more efficiently and enabling higher throughput data analysis. The resultant consensus genomes are assigned both PANGO lineage and variant status as defined by Public Health England. Samples from within individual runs are used to generate phylogenetic trees incorporating optional background samples as well as summaries of individual SNPs. As minoTour uses ARTIC pipelines, new primer schemes and pathogens can be added to allow minoTour to aid in real-time analysis of pathogens in the future.

**Availability and Implementation:** Source code and documentation is available at https://github.com/LooseLab/minotourapp.

**Supplementary information:** Supplementary data are available from

https://github.com/LooseLab/artic_minotour_analyses.

## Introduction

Oxford Nanopore Technologies (ONT) range of sequencers (MinION, GridION and Promethion) have transformed sequencing from a fixed to real-time process (Jain *et al*., 2016). By writing batches of sequenced reads to disk after DNA has finished translocating the pore, sequence data become available immediately, meaning data analysis can begin earlier, reducing the total time required to answer a specific question. The ongoing Covid-19 pandemic has provided a clear demonstration of the proposed benefits of real-time analysis of sequence data (Gardy and Loman, 2018; Quick *et al*., 2016), with questions such as lineage assignment and Variant of Concern/Variant under Investigation (VoC/VuI) status potentially being time sensitive (O’Toole *et al*., 2021). In theory this can be accelerated by incrementally analysing reads as they are created, a feature unique to ONT sequencers that we exploit here.

In our work as members of the COG-UK network (COVID-19 Genomics UK (COG-UK),2020), we generated thousands of SARS-CoV2 consensus sequences using ONT sequencers. To assess sequencing performance in real-time, we used minoTour, our real time analysis and monitoring system (Munro et. al., 2021) https://github.com/looselab/minotourapp), to track the performance of each sequencing run. Given that ONT flow cells, even using barcodes, can provide more data than required for an individual viral genome, it makes sense to stop sequencing once sufficient data are available for the analysis to be completed. Shorter sequencing runs preserve flow cell health, which can then be flushed and reused for other sequencing libraries and experiments. If efficiently implemented, this can reduce the cost per sample for sequencing. We therefore integrated these capabilities into minoTour.

The ARTIC Network (Tyson *et al*., 2020; Quick *et al*., 2017) (https://artic.network) provides comprehensive protocols for both wet lab and downstream state-of-the-art best practice informatics analysis for SARS-CoV2. The optimal ARTIC pipeline uses Nanopolish (Loman *et al*., 2015; Quick *et al*., 2017) for signal level analysis of Nanopore data during variant calling. As an alternative to Nanopolish, ARTIC provides medaka, a machine learning pipeline which only requires FASTQ data (https://github.com/nanoporetech/medaka). As signal level data are unavailable within minoTour, we integrated the ARTIC medaka workflow to enable real-time generation of consensus genomes in parallel with sequence data generation. This contrasts with other web based analysis platforms which either do not exploit the real-time features of the nanopore platform or do not have access to the sequence data themselves for further analysis (Ferguson *et al*., 2021; Bruno *et al*., 2021). The ARTIC network provides a tool, RAMPART, which can monitor a run over time and complete analysis for individual samples, but does not provide many of the other features shown here at this time (https://artic.network/rampart).

To enable automatic consensus generation minoTour uses a real-time predictive model for viral genome sequence coverage. Combining this with the ARTIC medaka pipeline, we can exploit the real-time features of Nanopore sequencing and demonstrate significant savings in run time, with resultant cost savings from flow cell reuse as well as faster identification of VoC/VuI’s in specific samples. MinoTour alerts the user through the Twitter API that a run can be stopped as well as providing visual feedback on the website that further sequencing may be a waste. The user can then use minoTour’s detailed breakdowns and visualisations of the ongoing sequencing run to make an informed decision on when to stop the run. Whilst this step could be automated, we have not implemented this as specific individual samples may be important enough that the user wishes for a run to continue, even with the reduction in useful data being generated.

We analyse the impact that stopping runs early may have on the generation of consensus genomes for samples by comparing the consensus sequences for 454 SARS-nCoV-2 samples assembled by both the medaka and nanopolish pipelines, investigating Phylogenetic Assignment of Named Global Outbreak (PANGO) lineage assignment, VoC/VuI assignment, as well as SNP calls for each sample at relevant timepoints during the run. We find a small loss of coverage from the last sample to complete on an individual flow cell. However, PANGO lineages and VoC/VuI assignments are all concordant within our 454 samples at all time points with both Nanopolish or Medaka based pipelines. Some SNP differences are observed, but these are predominantly ambiguous no calls (i.e an N at an individual position) rather than miss-calls.

## Implementation

### minoTour - A real-time LIMS system

MinoTour is free and open source software, written in python using the Django framework (see Munro et al). A separate command line client, minFQ (available via PyPi), uploads sequence data and sequencer metrics from the sequencing computer to the minoTour server. Once sequence data arrives, minoTour stores it in memory until it can be processed. Once processed and analysed, any sequence data are usually discarded, although they can optionally be stored in a database. Produced metrics are always stored in the database and visualised in the browser. MinoTour is compatible with Oxford Nanopore MinION and GridION sequencers and can process FASTQ files as generated by the Guppy base caller.

### Arctic Pipeline implementation

Our ARTIC based SARS-CoV2 python pipeline is executed as a Celery (https://docs.celeryproject.org/en/stable/) task, processing read batches pulled from minoTours memory cache. The pipeline is asynchronous, preventing blocking of any other analyses being performed. Reads are first filtered by length, with the minimum and maximum l calculated from the mean length of all amplicons in the scheme, plus/minus 50%, respectively. Reads are also filtered by the QC score assigned by Guppy, with only passing reads (Q score > 9, see Guppy release notes) being used in further analysis. The filtered reads are then mapped to an appropriate SARS-CoV-2 reference using minimap2 (Li, 2018). A numpy array of the length of the reference is created for each barcode as it is identified, and per base coverage is tracked using the mapped reads in real-time (van der Walt *et al*., 2011).

Coverage is tracked for each individual amplicon as defined by the primer scheme in use. Default parameters for triggering the analysis of a specific sample are at least 90% of the amplicons (completeness) covered at a median depth of at least 20x (coverage), but this can be changed by the user depending on requirements. Once this threshold has been passed, the accumulated mapped reads for that sample are passed to the ARTIC network medaka pipeline. The user can apply multiple coverage and completeness thresholds, re-triggering the analysis if more data becomes available. Varying primer schemes can be chosen, including custom schemes, simply by creating the appropriate primer scheme and reference files and uploading them to minoTour.

Once a consensus genome is available for a sample, minoTour uses pangolin (O’Toole *et al*., 2021) to assign a PANGO lineage from the most recent lineage classifications. After this step, the consensus sequence is compared with the current VoC/VuI definitions as defined by Public Health England (https://github.com/phe-genomics/variant_definitions) using the Aln2Type tool (https://github.com/connor-lab/aln2type). Both PANGO lineages and VoC/VuI designations are automatically updated daily by minoTour. Together, these generate a report for each sample (see Supplementary Figure 1C,D) and optionally the user can be notified if a VoC/VuI has been identified via the Twitter API. Finally, the sequences within each run are globally aligned using MAFFT (Katoh *et al*., 2002) and an illustrative tree generated using iQ-Tree (Minh *et al*., 2020) and visualised with figtree.js or ToyTree (Rambaut; Eaton, 2020). Additional background sequences can be included in these trees if desired and the distribution of SNPs within consensus sequences from the run compared with the reference are displayed in a snipit plot (https://github.com/aineniamh/snipit) (Supplementary Figure 1A). Links out from minoTour are provided to http://cov-lineages.org and http://outbreak.info for each specific variant identified as well as to the individual descriptors and data sources used to determine VoC/VuI classifications.

Results from the pipeline are maintained for historical record, with files stored on disk and metrics about the ARTIC sequencing experiment stored in a SQL database. These results are then visualised in the minoTour web server (discussed below). At any point, the user can choose to fire the ARTIC pipeline on one or all samples to investigate specific samples in more detail. Once a run has completed, automatically recognised by the fact that no further data are added to the flow cell within a fixed period of time, all analyses are automatically re-run to ensure maximum coverage for consensus generation. A retention policy for sequence data is set globally for the site and all read data can be automatically scrubbed from the server after consensus generation, if desired.

### Amplicon coverage prediction model

MinoTour predicts if individual samples are likely to result in an informative genome sequence, requiring no prior knowledge about sample numbers loaded on to an individual flow cell. MinoTour assumes the user is seeking minimal useful genome completeness (default 90% amplicons with at least 20X median “pass” read coverage). By using median depth, the impact of small insertions/deletions influencing our assessment of low coverage amplicons is reduced. The median coverage is only calculated for unique regions of each amplicon, excluding overlap to prevent artificial coverage inflation on schemes that contain amplicons with more than 50% overlap with neighbours. For making predictions, we assume that an ONT flow cell can generate at least 100,000 reads for each barcoded sample loaded and so project if each sample will reach minimal useful completeness. Our model’s algorithm is applied to each amplicon in the scheme (Equation 1). A sample is projected to finish if 90% of the amplicons have a predicted final coverage over the minimum required coverage (default 20X). All sequencing runs gather data for 1 hour before any of our strategies are used to ensure reasonable sampling of the loaded library.

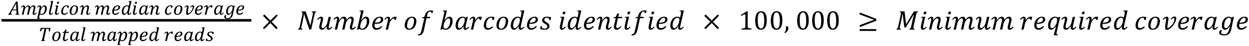

**Equation 1** - Used to predict if an amplicon will achieve the minimum required coverage during a run.

### ARTIC visualisations and reporting

Once the pipeline is underway, a new tab is added to the respective flow cell page, containing all data and visualisations pertaining to this ARTIC run (Supplementary Figure 1). The tab contains two sections of visualisations, with the top visualisations (Supplementary Figure 1A,B) showing the performance of all samples in the run, and the bottom section visualising a detailed performance for an individual sample. The visualised metrics include a bar chart displaying the read count for all barcoded samples, a bar chart displaying the mean read length of each sample in the run and the proportion of reads in the run that are unclassified vs. classified.

Below this a sortable and searchable summary table showing users metrics about each sample in the run, with average coverage, number of amplicons at >20X, >1X and 0X and basic statistics such as mean read length and read count displayed. If the sample has sufficient data to be run through the ARTIC pipeline, we display the assigned lineage and VoC/VoI status, and colour code the table row to indicate the sample has completed.

If a sample is selected to display more information, visualisations display a per base coverage across the sample genome, with plot bands displaying the primer schemes amplicon coordinates that the sequencing library was amplified with. Assigned PANGO lineage information is provided, with links out to further information describing each lineage (htpps://cov.lineages.org and https://outbreak.info). The VoC/VuI report generated by Aln2Type is visualised and the final status assigned displayed. A PDF report for each barcode and the overall run can be exported, showing all above metrics for each barcode (Supplementary File 1).

At the completion of a sequencing run, consensus sequence, pass and fail VCF files, BAM files and pangolin lineages can be downloaded. Optionally, these features can be disabled and minoTour will remove all files that may contain identifiable information from the server. By maintaining compatibility with standard ARTIC bioinformatics pipelines, this tool can be adapted to run any ARTIC compatible pathogen analysis simply by uploading the appropriate reference files.

### Post Run Genome Analysis

To investigate how manipulating run time affects results, we defined three timepoints of interest for a sample during a sequencing run. The full run time (FR), the run until time (RU) and the stop at time (SA). FR is defined as the time point at which the run completed with no intervention. RU is the point in a run where all samples our algorithm predicted would complete (90% completeness, 20X) had done so. Finally, SA is the point at which an individual sample in a run reached sufficient completeness and is automatically put through the ARTIC pipeline by minoTour, whilst the run continues.

To create consensus genomes from time points equivalent to our ARTIC pipeline and compare the results of both medaka and nanopolish we had to identify both the signal (FAST5) and FASTQ files equivalent to those minoTour would see. We mapped all reads from each barcode across all 13 reference ARTIC runs using minimap2 (Li, 2018) in file creation order. Using mosdepth (Pedersen and Quinlan, 2018) we determined coverage at each base across the reference genome and then median coverage for each amplicon using the same approach as in minoTour. This identifies the time points when sufficient data are available to trigger minoTour to analyse the genomes. The creation time point for the FASTQ file that results in sufficient coverage to meet the appropriate thresholds identifies the time in the sequencing run when analysis would occur. Using this method, we can identify the equivalent FAST5 files enabling us to analyse the data with both medaka and nanopolish. The code to generate this analysis is available from https://github.com/LooseLab/artic-minotour-analyses-scripts. For each of these timepoints, we generated consensus FASTA files to calculate genome recovery, defined as the proportion of non N positions in the final sequence. Whilst this is not a direct comparison to our completeness metric, it is a close approximation, as any base that has 20X coverage going into the ARTIC medaka pipeline will most likely be called as non N.

## Results and Discussion

### Amplicon coverage prediction model performance

The amplicon prediction model performed well across all runs (Figure 1A, R^2^=0.991). The model is conservative and underestimates true coverage, preventing waiting for genomes to complete which would never do so. Predicted genome recovery after an hour of data collection compares well with that observed at the calculated RU times for the 13 runs (Figure 1B). The strong correlation (R^2^=0.993) between predicted values and values actually recovered provides confidence in our algorithms performance. Comparing the RU stop time genome coverage with the FR coverage (Figure 1C) shows some small further benefits in coverage (R^2^=0.996). This is expected as continuing the run for longer allows the missing 10% of each genome to acquire some further coverage. However the longer a run continues the more this return diminishes, so stopping earlier accelerates time to answer as well as allowing the flow cell to be reused and so reduce cost. In addition, only the last genome to complete is typically affected. This can be seen more clearly (Figure 1D, R^2^=0.994) when filtering out those runs where no time is saved by our model. Genomes from these runs have the same RU and FR time, and so are identical by definition.

**Figure 1 -.**
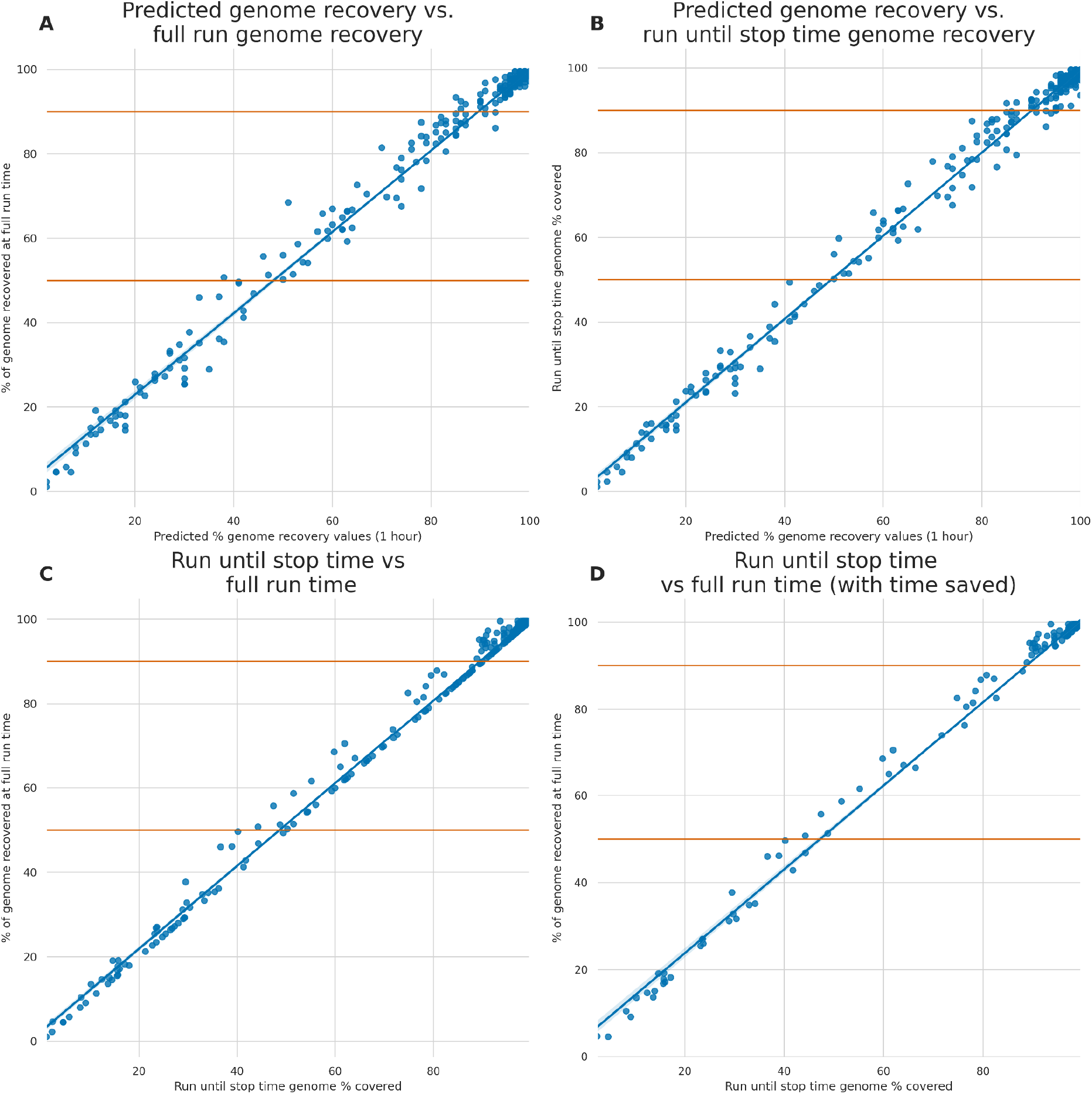
Genome recovery at time points throughout all 13 runs. **A)** The percentage of sample amplicons predicted to reach 20X coverage by our model using data from one hour of sequencing, compared to the percentage of the genome recovered (Non N bases) at FR by ARTICs medaka pipeline. **B)** The percentage of sample amplicons predicted to reach 20X coverage by our model using data from one hour of sequencing, compared to the percentage of the genome recovered (Non N bases) at the RU timepoint by ARTICs medaka pipeline. **C)** The percentage of the genome recovered (Non N bases) by ARTICs medaka pipeline at FR compared against the same at RU. **D)** The percentage of the genome recovered (Non N bases) by ARTICs medaka pipeline at FR compared against the same at RU, filtered for runs that would have saved time.

Our model predicts if a sample will generate sufficient data to provide useful information with enough accuracy to support a decision on whether or not to continue sequencing. There is a potential small loss in data as a consequence of reducing the sequencing time. We therefore quantify the consequences of this on time saved, lineage assignment and SNP calling below.

### Run until time saving

To quantify the time savings offered by our run until approach, we tracked metrics and predicted amplicon coverages per barcoded genome sample using minoTour for 13 sequencing runs. We plotted the time we would stop a run according to our model and the full length of the run (Figure 2A). Runs 9 through 13 were actively monitored with minoTour and manually stopped at earlier run times in response to the model predictions. Time savings using this approach are dependent on the sample composition, but are often significant (for example, Run 4, Figure 2A). Time savings are greater in runs with fewer samples as each has relatively more sequencing capacity available as can be seen in Figure 2B-N.

**Figure 2 -.**
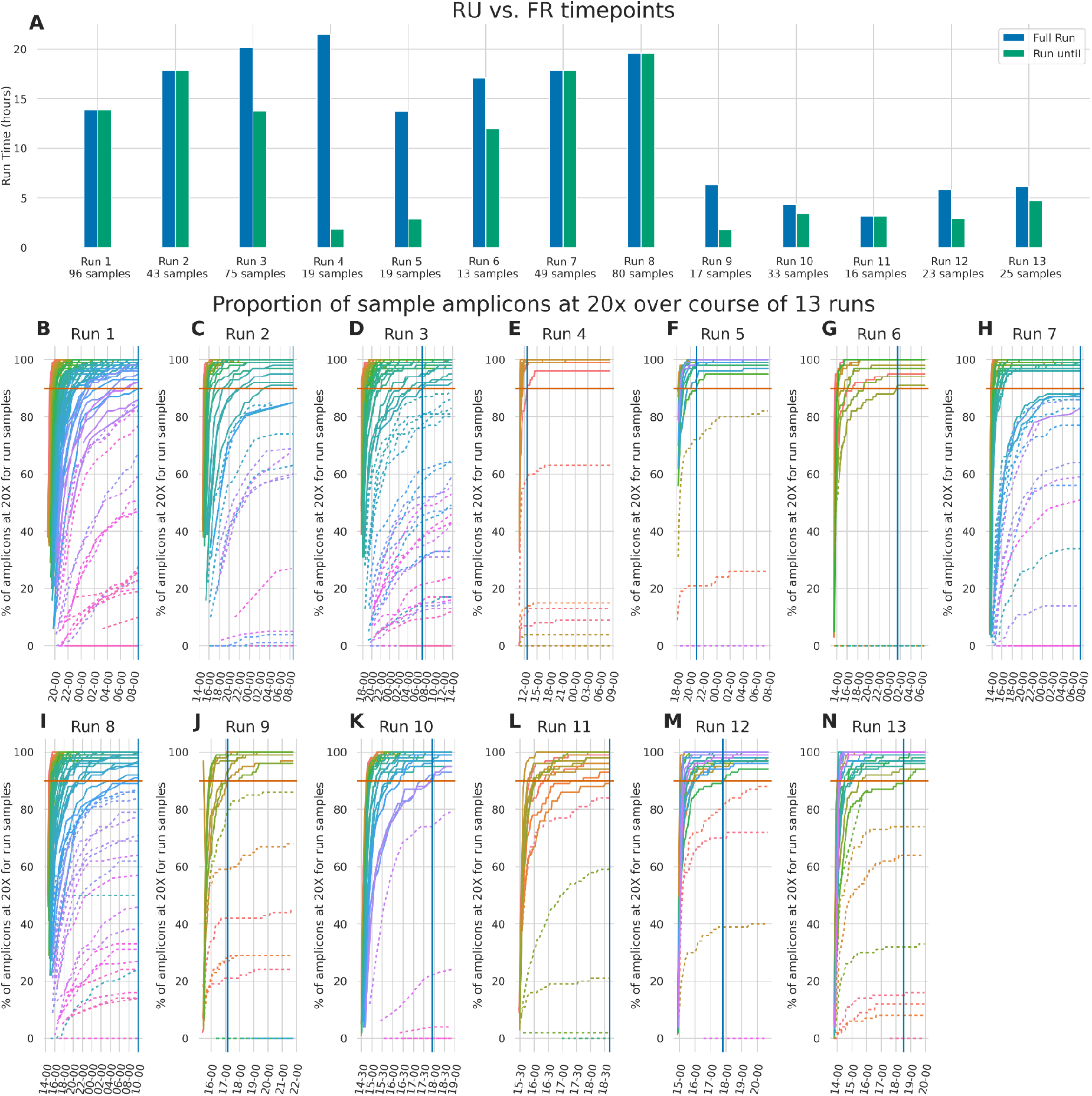
**A)** The FR time point plotted alongside the RU time point as hours since the run started. **B-N)** Samples across the course of 13 runs showing the percentage of amplicons at 20x. Barcodes that we project to finish are displayed with solid lines, whilst barcodes we project not to finish are dashed. 90% (Our threshold for firing) is marked on each plot. Once all barcodes that are projected to finish cross the 90% threshold, we would instruct MinKNOW to stop the run. This time is marked by a solid blue vertical line.

### Genome recovery

We next investigated the impact of stopping a run early on genome recovery. The number of genomes predicted to finish after one hour of sequencing with 20x median coverage across 90% of amplicons is very similar to those that actually do so (331/334 genomes predicted to finish do so, whilst 3/174 genomes not predicted to finish do, see Figure 3A). Where we predict more samples finish than will do so (e.g Run 2, Run 7, Run 11) no time savings are possible as minoTour will wait for these samples to complete before recommending stopping the run. In all cases this is caused by samples reaching a completeness just short of the 90% threshold but theoretically being able to complete given more reads. These nuances are not yet fully captured in our model and so users can fine tune these parameters as they need.

**Figure 3 -.**
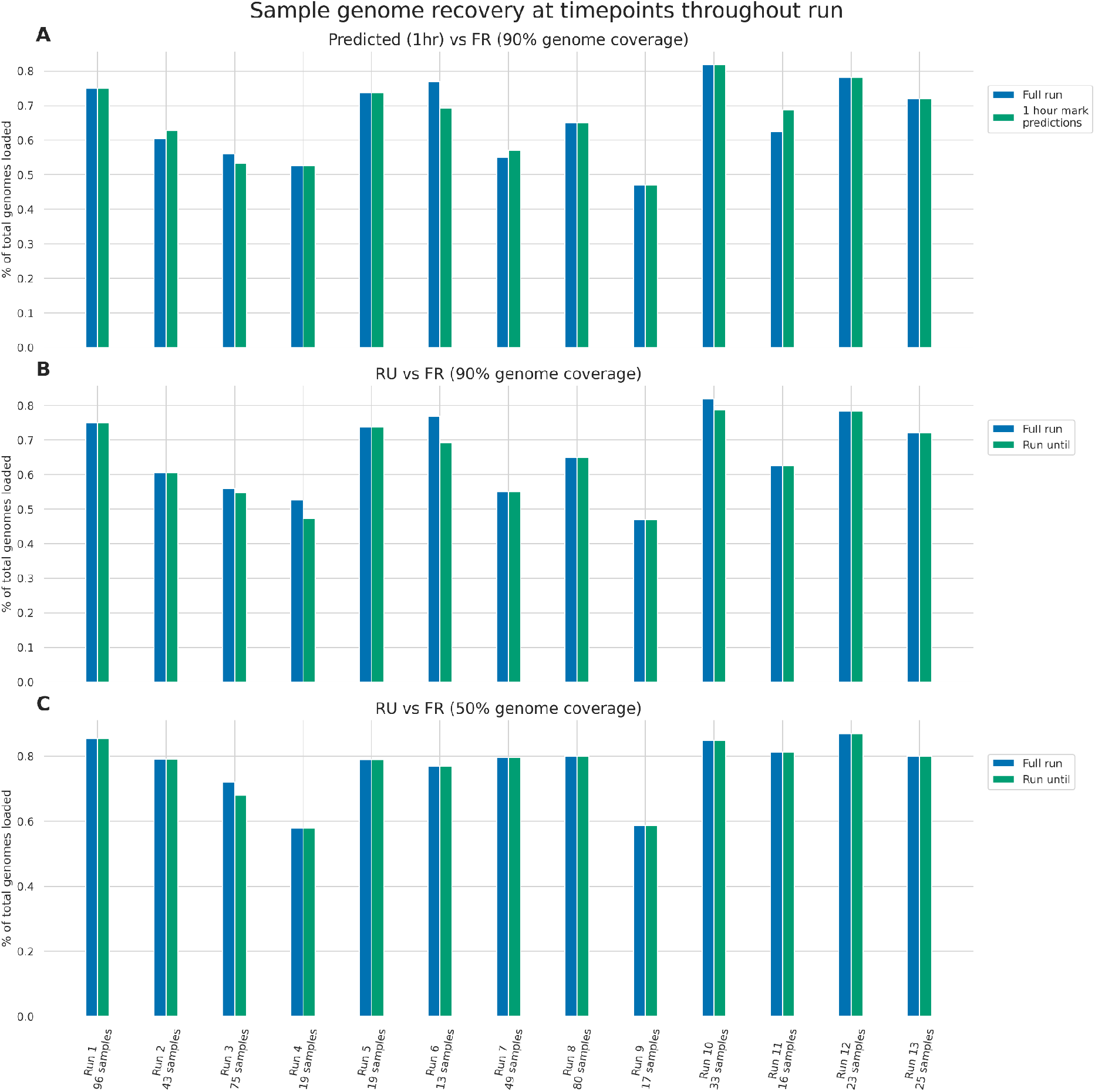
Genome completeness across 13 runs. **A)** Genomes predicted with one hours data to achieve 20x median coverage across 90% of amplicons compared with genomes that were recovered at 90% completeness (% non N bases) at FR, as a percentage of total genomes loaded onto flow cells. **B)** Genomes with 90% of amplicons covered at 20x median coverage at RU compared with genomes recovered at 90% genome completeness (% non N bases) at FR, as a percentage of total genomes loaded onto flow cells. **C)** Genomes recovered at 50% of amplicons covered at 20x at RU stop time compared with genomes recovered at 50% genome completeness (% non N bases) at FR as a percentage of total genomes loaded onto flow cells.

Figure 3B shows samples recovered at RU vs FR, wth 90% genome coverage. In almost all cases, we capture as much genomic data when stopping the run early as we would have by continuing sequencing. In only one case, Run 4, two samples failed to reach the maximum possible coverage. This was an artifact of marginal gains in sample coverage from allowing the run to continue for longer. Of course, if the user varies the minimum completeness of the genome required, the behaviour changes. Figure 3C illustrates genome recovery when specifying only 50% of amplicons are covered at a median depth of 20x. Only one run (Run 3) had a difference between samples recovered at RU and the FR timepoints.

### Lineage and SNP calling analysis

#### Lineage assignment to consensus genomes

Across all 13 sequencing runs, a total of 508 SARS-CoV-2 genomes were sequenced (including negative and positive controls). Of these, 456 produced genomes with both medaka and nanopolish ARTIC pipelines at FR, 454 produced genomes at RU and 335 genomes were produced at SA. The two additional genomes at the FR time are both extremely low completeness genomes (only 1% of the genome has consensus sequence) that failed to call at the RU time. Across all time points for any given sample in any run, we observe complete concordance in lineage assignment between either medaka or nanopolish analysed genomes (Supplementary File 2). Any loss of data seen by stopping sequencing early did not impact PANGO lineage assignment in a SARS-CoV-2 sequencing run. We note that these sequences are predominantly from the B.1.1.7 lineage due to the timeframe in which they were collected, but given our observations on SNP calling below do not envisage this being an issue.

#### Comparing SNPs between medaka and nanopolish consensus genomes

We compared nanopolish and medaka consensus genome sequences for each genome in our data set (1,245 genomes from 456 unique samples) constructed from all three timepoints (FR, RU and SA) (Figure 4A-C). The SNPs were called using nextclade (https://clades.nextstrain.org) with the output data available in Supplementary Files 3 and 4. From all 13 runs, 456 unique samples generated a consensus genome at FR with both medaka and nanopolish (Figure 4A). 341 of these samples were identical between the two pipelines. The remaining 115 genomes have at least one difference in the consensus sequence. The majority of these are where either medaka or nanopolish are unable to confidently call a site and so assigns an ambiguous base (N). Of more concern, there are some sites which are incorrectly assigned as a reference call by medaka. On closer inspection, the majority of these are for one single site in the genome at position 28,111 (Figure 4I). At the RU timepoint, a similar pattern is observed with a small increase in the number of ambiguous (N) sites, most likely a consequence of the slightly lower coverage data available (Figure 4B). At the most extreme, the SA time, the number of ambiguous sites increases further (Figure 4C). It is worth noting that on any multiplexed run, only one sample ever finishes at this time point. All other samples, by definition, are complete at the RU timepoint and so are expected to be of higher quality. Thus the majority of SNP differences between medaka and nanopolish are differences in ambiguous calls.

**Figure 4 -.**
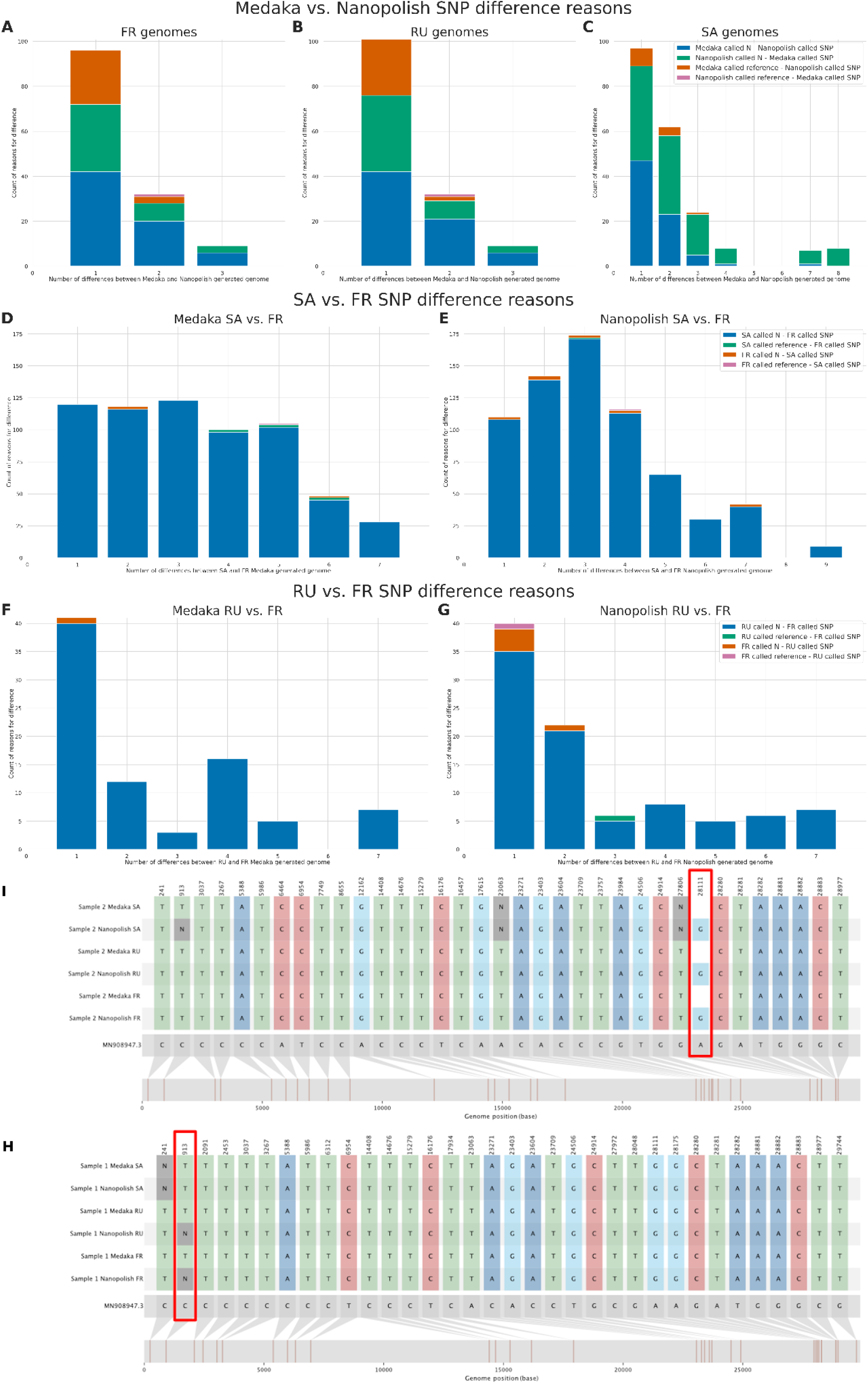
Comparison of SNPs called from consensus genomes generated by ARTICs nanopolish and medaka pipelines. **A-C)** Comparison of sample SNPs from consensus genomes generated by medaka and nanopolish, at FR, RU and SA. Samples are split by the number of differing SNP calls between them, with the reason for differences colour coded into bars. **D-G)** Comparison of sample SNPs at FR and SA, and FR and RU, for consensuses generated by either medaka or nanopolish. Samples are split by the number of differing SNP calls between them, and the reason for the difference is colour coded into bars. **H)**. SNIPIT plot of an example pair of consensus genomes, showing the nanopolish pipeline switch from a SNP to an N at position 913 with more data. **I)**. SNIPIT plot showing an example pair of consensus genomes with nanopolish calling a SNP at position 28111 but medaka calling reference.

#### Comparing SNPs between timepoints with medaka and nanopolish

To determine if stopping a run early was detrimental to calling SNPs using either pipeline, we compared SNP calls between the FR and RU, and FR and SA timepoints for genomes produced by both medaka and nanopolish (Figure 4D-G). As expected, the vast majority of these differences are conservative ambiguous (N) calls when less sequencing data are available for consensus generation. Nanopolish appears more conservative in this regard resulting in more ambiguous calls than medaka. We find one site, 913, for which Nanopolish rarely can call a SNP at lower coverage, but changes to an ambiguous call at higher depth

SNIPIT plots illustrating the differences between genomes (Figure 4H,I) provide examples of the various SNP artifacts caused by changes in coverage and pipeline. Of concern are those positions where a reference site is called when a SNP is likely present (Figure 4I). This is most common at position 28,111 (53/69 occurrences) and rare elsewhere in the genome. Overall, we find that medaka is sufficient for variant calling and lineage assignment, but in our workflows we routinely ran both pipelines for confirmation. Handling signal data in minoTour is not currently feasible, excluding the nanopolish pipeline from our analyses.

## Conclusions

The use of real-time sequencing has been proposed as a key component of pathogen surveillance during disease outbreaks (Gardy and Loman, 2018). The SARS-CoV2 pandemic has seen the development of new tools and enhancement of existing approaches for collating, sharing and analysing sequence data at local, national and international scales (Nicholls *et al*., 2021; COVID-19 Genomics UK (COG-UK),2020; Shu and McCauley, 2017; Hadfield *et al*., 2018). Portable sequencers, such as those developed by Oxford Nanopore, can be readily deployed in the field and samples can be sequenced anywhere. Excellent analysis pipelines are available to generate consensus genomes, but even though well documented, can be beyond some users to apply (Tyson *et al*., 2020). Here we show how our existing Nanopore toolset, minoTour, can be extended to include real-time analysis using the best available practices for SARS-CoV2 sequencing.

By formalising a model predicting likely coverage per sample, we enable the user to predict the performance of a sequencing run and better determine when to stop sequencing. This approach leads to cost and time savings in many instances. Benefits of centralising analysis include standardisation of analysis pipelines as well as the opportunity to incorporate additional reporting such as lineage and VoC/VuI assignment as well as linking out to useful further resources. The generation of consensus sequence and lineage assignment happens during the sequencing run, enabling the potential for rapid feedback of results if required.

MinoTour can be installed on a single laptop/computer running sequencing or can be run as a central hub with data uploaded by multiple devices concurrently. Docker and standard Django installation instructions are available. In our experience, a laptop powerful enough to run a nanopore sequencer with real time base-calling is sufficient to also run minoTour. The exception to this is the minIT/MK1C device which cannot be configured to run minoTour at present. However, these can be used with the minFQ client for data upload. Thus minoTour can serve as a hub for data collection from multiple sites. We have engineered the ARTIC pipeline in minoTour such that any ARTIC compatible primer scheme can be incorporated and used for analysis. As proof of principle we have tested this with multiple SARS-CoV2 schemes as well as using historical data from studies of Ebola (Quick *et al*., 2016). A single instance of minoTour can run multiple ARTIC pipelines tailored for specific pathogens, although additional features such as PANGO lineage assignment and variant detection pipelines are pathogen specific. Uniquely, and in contrast to other tools, minoTour exploits almost all the real-time features of the Nanopore sequencing platform for maximising ARTIC sequencing speed and simplifying analysis.

## Acknowledgements

Thanks to John Tyson and members of the COG-UK Consortium for helpful comments on early prototypes of the code presented here.

