## Supplementary File 1 for "Real-time monitoring and analysis of SARS-CoV-2 nanopore sequencing with minoTour": 103_artic_report.pdf

### Report for artic output of FAQ01023_CV197_25_M1 generated 8th September 2021 14:18

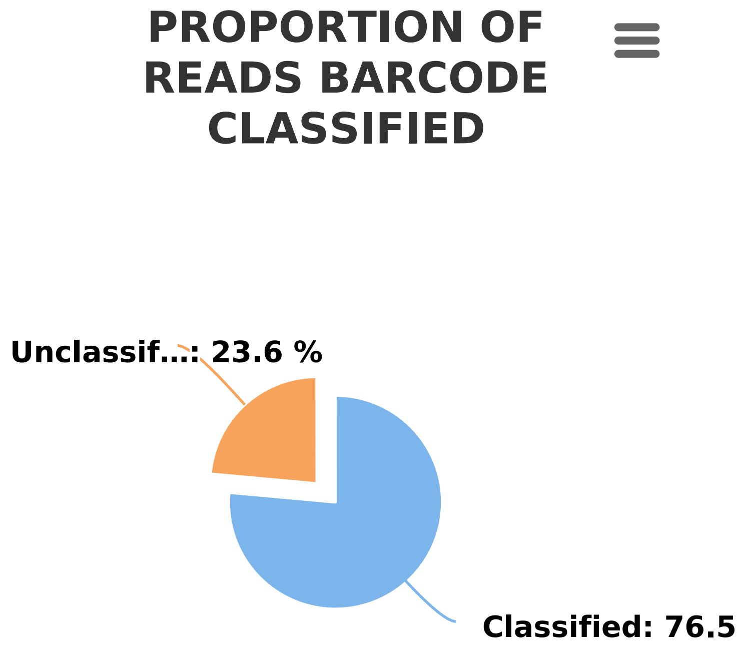


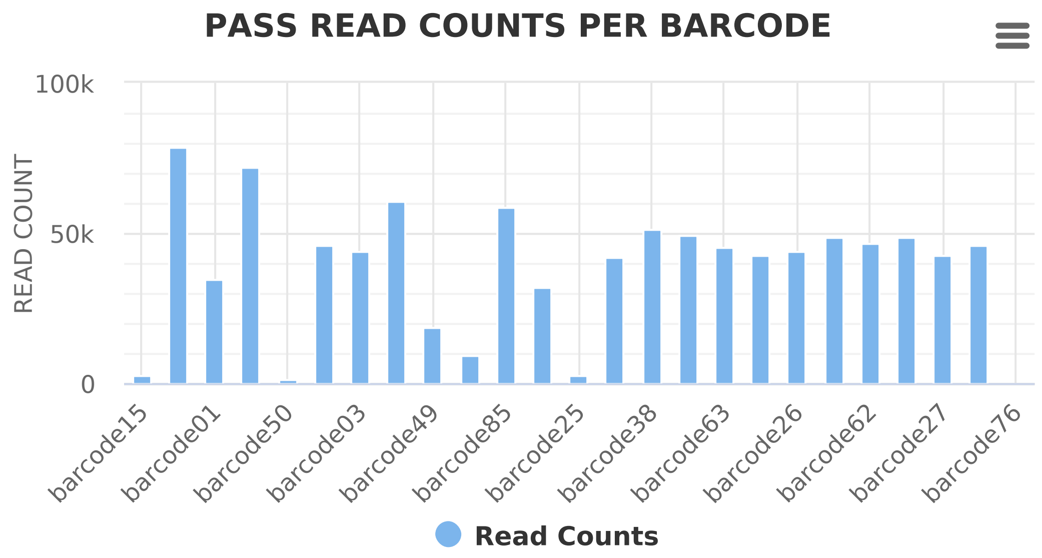


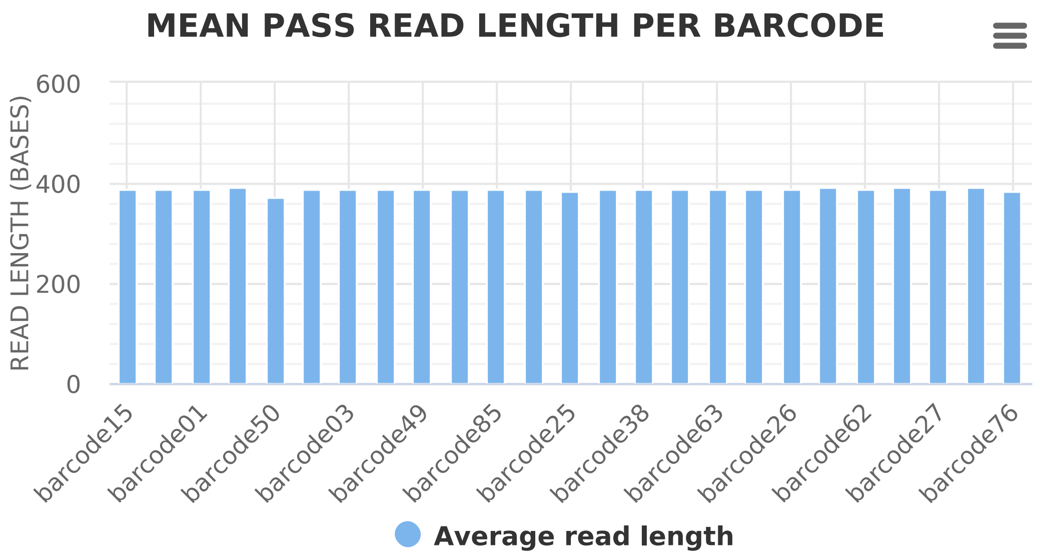


| Barcode Name | Chromosome | Read Len | Read Count | Yield | Coverage | # Success Amplicons | # Partial Amplicons | # Failed Amplicons | Lineage | VoC found |
|---|---|---|---|---|---|---|---|---|---|---|
| barcode01 | MN908947.3 | 388 | 34,489 | 13,393,773 | 438.88 | 76 | 4 | 19 | Currently unknown | Not Tested |
| barcode02 | MN908947.3 | 388 | 49,127 | 19,051,766 | 624.2 | 99 | 0 | 0 | B.1.1.7 | 0 VOC-20DEC-01 Name: phe-label, dtype: object |
| barcode03 | MN908947.3 | 389 | 44,022 | 17,106,691 | 560.4 | 94 | 5 | 0 | Currently unknown | Not Tested |
| barcode13 | MN908947.3 | 388 | 32,321 | 12,537,233 | 410.86 | 88 | 9 | 2 | Currently unknown | Not Tested |
| barcode14 | MN908947.3 | 391 | 45,820 | 17,913,802 | 587.57 | 99 | 0 | 0 | B.1.1.7 | 0 VOC-20DEC-01 Name: phe-label, dtype: object |
| barcode15 | MN908947.3 | 388 | 2,415 | 936,286 | 30.72 | 17 | 5 | 77 | Currently unknown | Not Tested |
| barcode25 | MN908947.3 | 386 | 2,348 | 906,274 | 29.77 | 8 | 2 | 89 | Currently unknown | Not Tested |
| barcode26 | MN908947.3 | 390 | 44,063 | 17,173,790 | 563.16 | 99 | 0 | 0 | B.1.1.7 | 0 VOC-20DEC-01 Name: phe-label, dtype: object |

| Barcode Name | Chromosome | Read Len | Read Count | Yield | Coverage | # Success Amplicons | # Partial Amplicons | # Failed Amplicons | Lineage | VoC found |
|---|---|---|---|---|---|---|---|---|---|---|
| barcode27 | MN908947.3 | 387 | 42,641 | 16,520,274 | 541.29 | 96 | 3 | 0 | Currently unknown | Not Tested |
| barcode37 | MN908947.3 | 387 | 46,152 | 17,881,875 | 585.97 | 91 | 5 | 3 | Currently unknown | Not Tested |
| barcode38 | MN908947.3 | 388 | 51,030 | 19,810,123 | 649.61 | 98 | 0 | 1 | B.1.1.7 | 0 VOC-20DEC-01 Name: phe-label, dtype: object |
| barcode39 | MN908947.3 | 388 | 42,319 | 16,437,763 | 538.77 | 95 | 4 | 0 | B.1.1.7 | 0 VOC-20DEC-01 Name: phe-label, dtype: object |
| barcode49 | MN908947.3 | 387 | 18,946 | 7,325,827 | 240.15 | 63 | 7 | 29 | Currently unknown | Not Tested |
| barcode50 | MN908947.3 | 374 | 1,506 | 563,440 | 18.46 | 16 | 3 | 80 | Currently unknown | Not Tested |
| barcode51 | MN908947.3 | 391 | 72,218 | 28,269,815 | 927.41 | 99 | 0 | 0 | B.1.1.7 | 0 VOC-20DEC-01 Name: phe-label, dtype: object |
| barcode61 | MN908947.3 | 388 | 9,602 | 3,724,731 | 122.27 | 34 | 4 | 61 | Currently unknown | Not Tested |
| barcode62 | MN908947.3 | 390 | 46,897 | 18,282,670 | 599.62 | 99 | 0 | 0 | B.1.1.7 | 0 VOC-20DEC-01 Name: phe-label, dtype: object |
| barcode63 | MN908947.3 | 390 | 45,320 | 17,654,226 | 579.03 | 99 | 0 | 0 | B.1.1.7 | 0 VOC-20DEC-01 Name: phe-label, dtype: object |
| barcode73 | MN908947.3 | 391 | 48,380 | 18,893,415 | 619.56 | 99 | 0 | 0 | B.1.1.7 | 0 VOC-20DEC-01 1 E484K Name: phe-label, dtype: object |
| barcode74 | MN908947.3 | 390 | 42,751 | 16,659,804 | 546.41 | 99 | 0 | 0 | B.1.1.7 | 0 VOC-20DEC-01 Name: phe-label, dtype: object |
| barcode75 | MN908947.3 | 389 | 60,771 | 23,647,077 | 774.72 | 97 | 2 | 0 | B.1.1.7 | 0 VOC-20DEC-01 Name: phe-label, dtype: object |
| barcode76 | MN908947.3 | 384 | 1 | 384 | 0.01 | 0 | 1 | 98 | Currently unknown | Not Tested |
| barcode85 | MN908947.3 | 389 | 58,861 | 22,924,435 | 751.68 | 99 | 0 | 0 | B.1.1.7 | 0 VOC-20DEC-01 Name: phe-label, dtype: object |
| barcode86 | MN908947.3 | 392 | 48,633 | 19,063,992 | 625.28 | 99 | 0 | 0 | B.1.1.7 | 0 VOC-20DEC-01 Name: phe-label, dtype: object |
| barcode87 | MN908947.3 | 389 | 78,605 | 30,562,375 | 1,002.43 | 99 | 0 | 0 | B.1.1.7 | 0 VOC-20DEC-01 Name: phe-label, dtype: object |
| unclassified | MN908947.3 | 369 | 298,621 | 110,329,619 | 3,595.19 | 99 | 0 | 0 | Currently unknown | Not Tested |

### Artic generated SNIPIT plot and tree for FAQ01023_CV197_25_M1

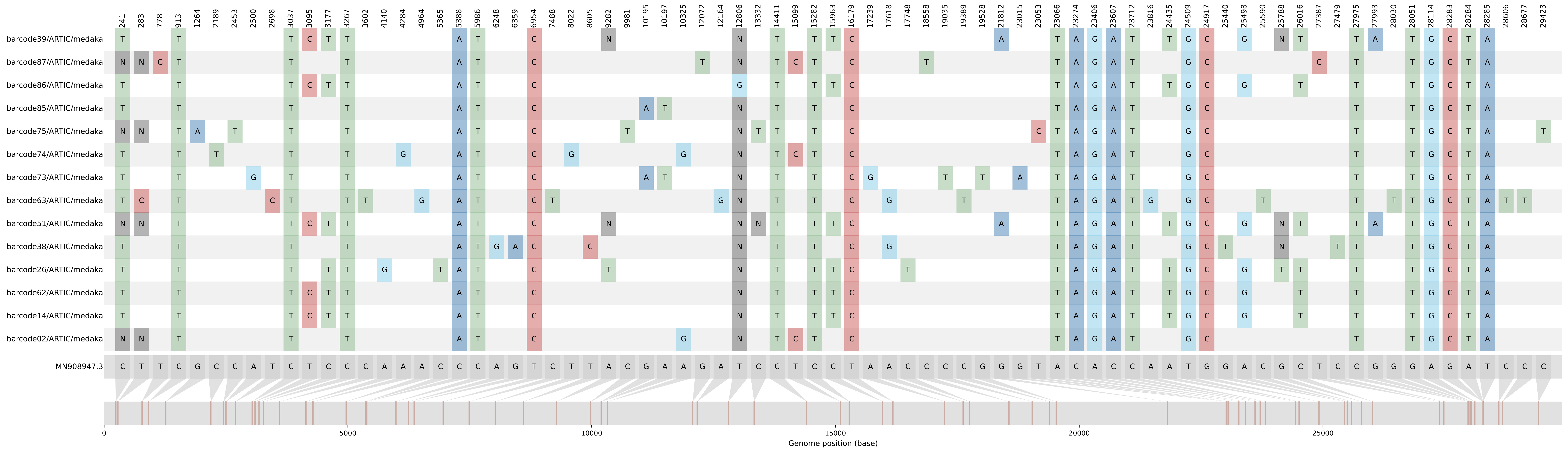

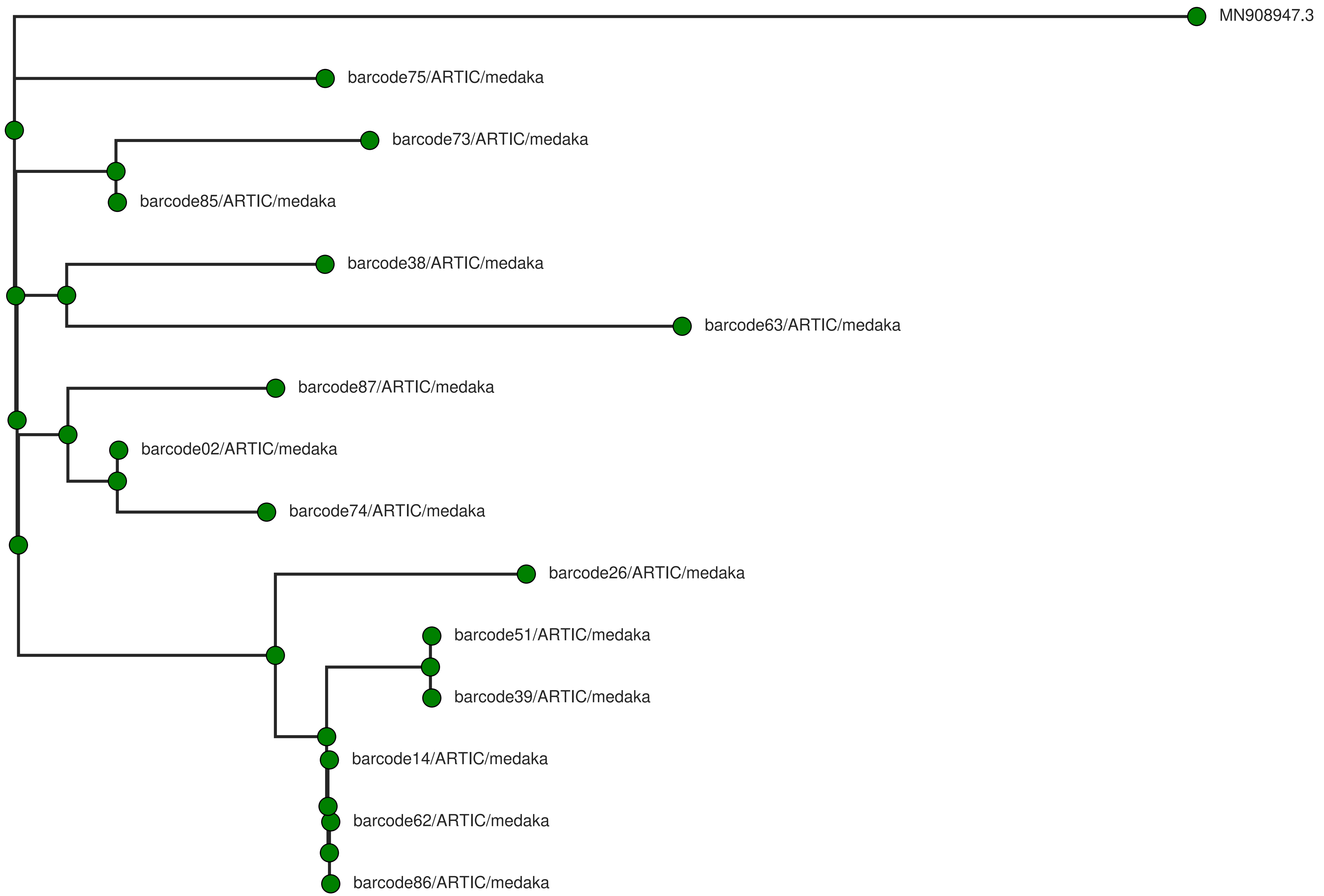
MN908947.3
barcode75/ARTIC/medaka
barcode73/ARTIC/medaka
barcode85/ARTIC/medaka
barcode38/ARTIC/medaka
barcode63/ARTIC/medaka
barcode87/ARTIC/medaka
barcode02/ARTIC/medaka
barcode74/ARTIC/medaka
barcode26/ARTIC/medaka
barcode51/ARTIC/medaka
barcode39/ARTIC/medaka
barcode14/ARTIC/medaka
barcode62/ARTIC/medaka
barcode86/ARTIC/medaka
