## Supplementary File 1 for "Real-time monitoring and analysis of SARS-CoV-2 nanopore sequencing with minoTour": 103_barcode02_artic_report.pdf

### Report for artic output of barcode02 generated 8th September 2021 14:18

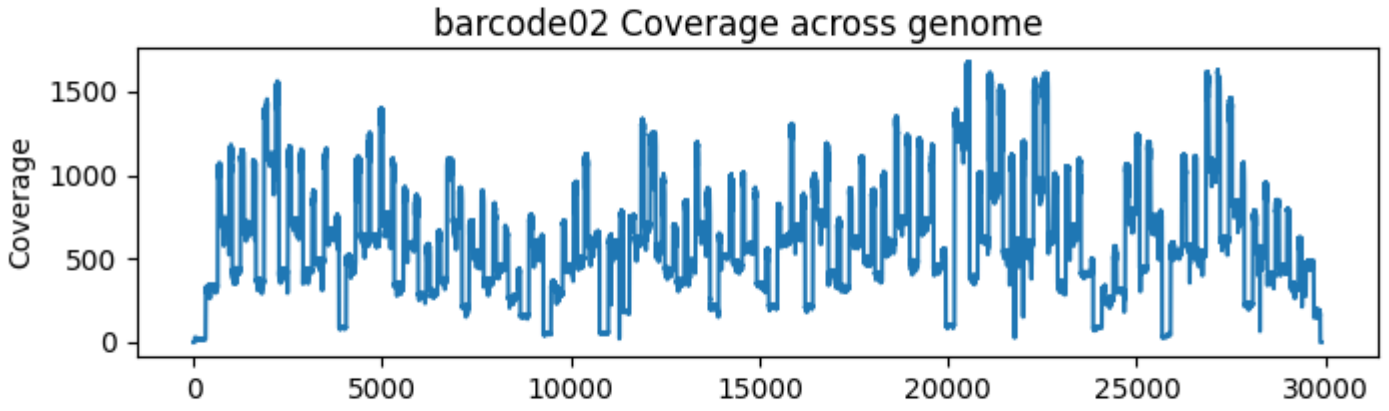

### Artic variant summaries for barcode02

#### Variant of Concern Report for barcode02/ARTIC/medaka

If a genome is analysed, this will provide a report for the variant or variants found based on current PHE Variant Definitions. This analysis will report all possible VoCs and caution should be taken in interpretation of low coverage/quality genomes.

Reported VuI/VoCs:

| sample_id | phe-label | unique-id | status | mutation-ref-calls | mutation-mixed-calls | mutation-calls | indel-ref-calls | indel-calls | no-calls | no-calls-deletion |
|---|---|---|---|---|---|---|---|---|---|---|
| barcode02/ARTIC/medaka | VOC-20DEC-01 | denture-daughter | confirmed | 0 | 0 | 13 | 0 | 2 | 0 | 0 |

Observed mutations:

| | 0 | 1 | 2 | 3 | 4 | 5 | 6 | 7 | 8 | 9 | 10 |
|---|---|---|---|---|---|---|---|---|---|---|---|
| **sampleID** | barcode02/ ARTIC/ medaka | barcode02/ ARTIC/ medaka | barcode02/ ARTIC/ medaka | barcode02/ ARTIC/ medaka | barcode02/ ARTIC/ medaka | barcode02/ ARTIC/ medaka | barcode02/ ARTIC/ medaka | barcode02/ ARTIC/ medaka | barcode02/ ARTIC/ medaka | barcode02/ ARTIC/ medaka | barcode02/ ARTIC/ medaka |
| **type** | snp | snp | snp | snp | snp | snp | snp | del | snp | snp | snp |
| **reference-base** | C | C | C | C | C | T | A | GTCTGGTTTT | C | T | C |
| **variant-base** | T | T | T | A | T | C | G | G | T | C | T |
| **var-length** | 1 | 1 | 1 | 1 | 1 | 1 | 1 | 10 | 1 | 1 | 1 |
| **one-based-reference-position** | 913 | 3037 | 3267 | 5388 | 5986 | 6954 | 10323 | 11287 | 14408 | 15096 | 15279 |
| **iupac-variant-bases** | T | T | T | A | T | C | G | G | T | C | T |

| | 11 | 12 | 13 | 14 | 15 | 16 | 17 | 18 | 19 | 20 | 21 |
|---|---|---|---|---|---|---|---|---|---|---|---|
| **sampleID** | barcode02/ ARTIC/ medaka | barcode02/ ARTIC/ medaka | barcode02/ ARTIC/ medaka | barcode02/ ARTIC/ medaka | barcode02/ ARTIC/ medaka | barcode02/ ARTIC/ medaka | barcode02/ ARTIC/ medaka | barcode02/ ARTIC/ medaka | barcode02/ ARTIC/ medaka | barcode02/ ARTIC/ medaka | barcode02 ARTIC/ medaka |
| **type** | snp | del | del | snp | snp | snp | snp | snp | snp | snp | snp |
| **reference-base** | T | ATACATG | TTTA | A | C | A | C | C | T | G | C |
| **variant-base** | C | A | T | T | A | G | A | T | G | C | T |
| **var-length** | 1 | 7 | 4 | 1 | 1 | 1 | 1 | 1 | 1 | 1 | 1 |
| **one-based-reference-position** | 16176 | 21764 | 21990 | 23063 | 23271 | 23403 | 23604 | 23709 | 24506 | 24914 | 27972 |
| **iupac-variant-bases** | C | A | T | T | A | G | A | T | G | C | T |

Sample_ID: barcode02/ARTIC/medaka
PANGO:B.1.1.7
nextstrain:N501Y.V1

Variant Status: confirmed

PHE-Label: VOC-20DEC-01
WHO Label: Alpha
Alternate Names: VOC202012/01, UK variant, Kent variant, VOC1,

*Description:*
This variant became widespread in the UK in the Winter of 2021 and is characterised by increased transmissibility.
*Information Sources* Source_1 Source_2

Variant Calls Detected:

| Position | gene | protein | ref | variant | sample call | type | status |
|---|---|---|---|---|---|---|---|
| 3267 | ORF1ab | nsp3 | C | T | T | SNP | detect |
| 5388 | ORF1ab | nsp3 | C | A | A | SNP | detect |
| 6954 | ORF1ab | nsp3 | T | C | C | SNP | detect |
| 21764 | S | surface glycoprotein | ATACATG | A | A | deletion | detect |
| 21990 | S | surface glycoprotein | TTTA | T | T | deletion | detect |
| 23063 | S | surface glycoprotein | A | T | T | SNP | detect |
| 23271 | S | surface glycoprotein | C | A | A | SNP | detect |
| 23604 | S | surface glycoprotein | C | A | A | SNP | detect |
| 23709 | S | surface glycoprotein | C | T | T | SNP | detect |
| 24506 | S | surface glycoprotein | T | G | G | SNP | detect |
| 24914 | S | surface glycoprotein | G | C | C | SNP | detect |
| 27972 | ORF8 | ORF8 protein | C | T | T | SNP | detect |
| 28048 | ORF8 | ORF8 protein | G | T | T | SNP | detect |
| 28111 | ORF8 | ORF8 protein | A | G | G | SNP | detect |
| 28280 | N | nucleocapsid phosphoprotein | GAT | CTA | CTA | MNP | detect |

One based position - notes on sites detected in this sample.

The observed calls were:

| |
|---|
| mutation_ref_calls: 0 |
| indel_ref_calls: 0 |
| mutation_calls: 13 |
| mutation_mixed_calls: 0 |
| indel_calls: 2 |
| no_calls: 0 |
| no_call_deletion: 0 |

Classification rules:

confirmed

mutations_required: 13

indels_required: 0

allowed_wildtype: 0

probable

mutations_required: 5

indels_required: 0

allowed_wildtype: 0

low_qc

mutations_required: 0

indels_required: 0

allowed_wildtype: 0

*Acknowledgements:*
*Curators:*Natalie Groves Ulf Schaefer Nick Loman

The complete list of variants of concern scanned included:
VUI-21FEB-04 VOC-20DEC-02 VUI-21FEB-01 VUI-21MAR-01 VUI-21MAY-01 VOC-20DEC-01 VOC-21APR-02 VUI-21FEB-03 VUI-21APR-01 E484K VUI-21JUL-01 VUI-21JAN-01 VUI-21MAR-02 VUI-21MAY-02 VUI-21JUN-01 VOC-21JAN-02 VUI-21APR-03 VOC-21FEB-02
