## Supplementary File 1 for "Real-time monitoring and analysis of SARS-CoV-2 nanopore sequencing with minoTour": 103_barcode50_artic_report.pdf

### Report for artic output of barcode50 generated 8th September 2021 14:18

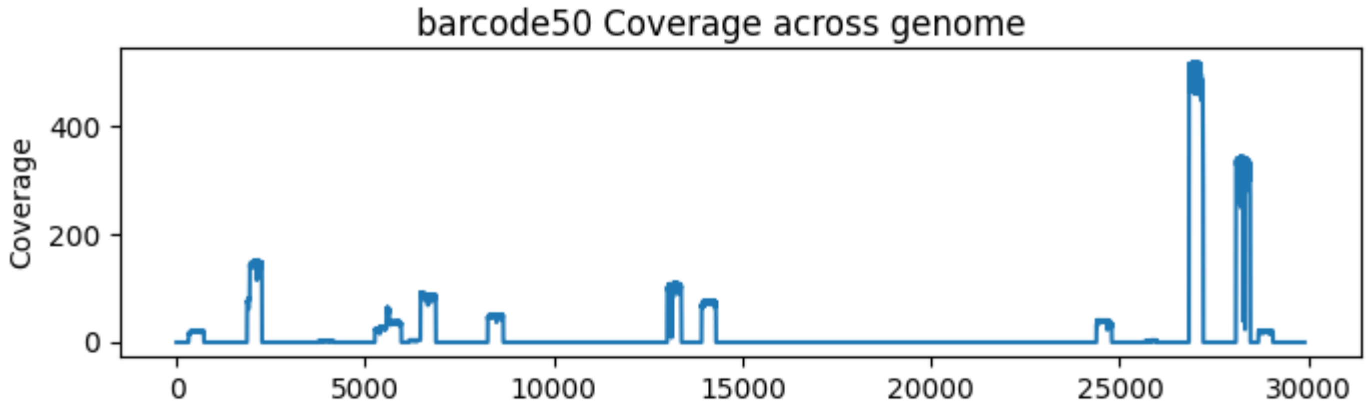

### Artic variant summaries for barcode50

#### Variant of Concern Reports not available.

If a genome is complete and has a Variant of Concern lineage assigned to it, summary reports will be provided here based on current PHE Variant Definitions.
