## Supplementary File 1 for "Real-time monitoring and analysis of SARS-CoV-2 nanopore sequencing with minoTour": 103_barcode73_artic_report.pdf

Report for artic output of barcode73 generated 8th September 2021 14:18

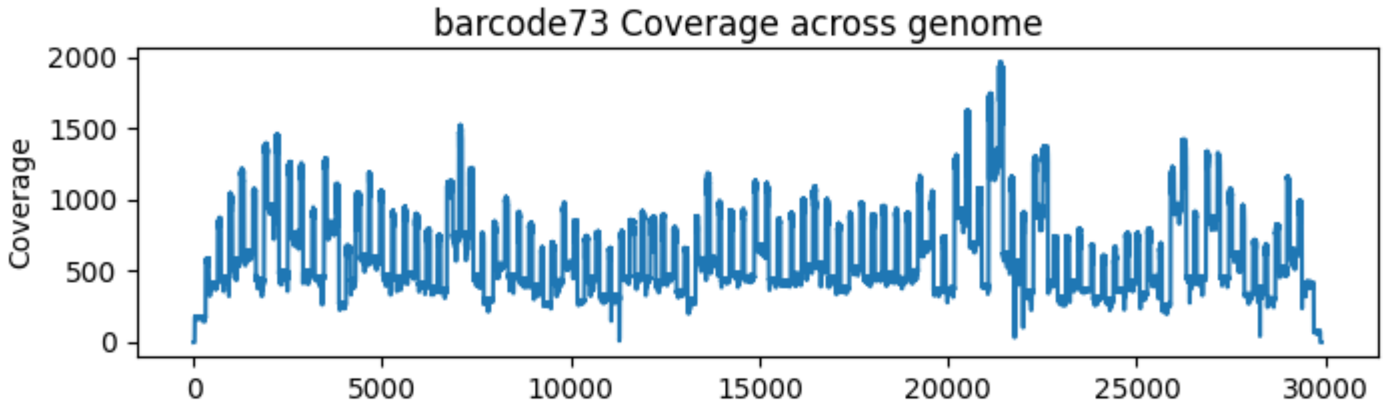

### Artic variant summaries for barcode73

#### Variant of Concern Report for barcode73/ARTIC/medaka

If a genome is analysed, this will provide a report for the variant or variants found based on current PHE Variant Definitions. This analysis will report all possible VoCs and caution should be taken in interpretation of low coverage/quality genomes.

Reported VuI/VoCs:

| sample_id | phe-label | unique-id | status | mutation-ref-calls | mutation-mixed-calls | mutation-calls | indel-ref-calls | indel-calls | no-calls | no-calls-deletion |
|---|---|---|---|---|---|---|---|---|---|---|
| barcode73/ARTIC/medaka | VOC-20DEC-01 | denture-daughter | confirmed | 0 | 0 | 13 | 0 | 2 | 0 | 0 |
| barcode73/ARTIC/medaka | E484K | harbor-sprite | confirmed | 0 | 0 | 1 | 0 | 0 | 0 | 0 |

Observed mutations:

| | 0 | 1 | 2 | 3 | 4 | 5 | 6 | 7 | 8 | 9 | 10 |
|---|---|---|---|---|---|---|---|---|---|---|---|
| **sampleID** | barcode73/ARTIC/medaka | barcode73/ARTIC/medaka | barcode73/ARTIC/medaka | barcode73/ARTIC/medaka | barcode73/ARTIC/medaka | barcode73/ARTIC/medaka | barcode73/ARTIC/medaka | barcode73/ARTIC/medaka | barcode73/ARTIC/medaka | barcode73/ARTIC/medaka | barcode73/ARTIC/medaka |
| **type** | snp | snp | snp | snp | snp | snp | snp | snp | snp | snp | del |
| **reference-base** | C | C | A | C | C | C | C | T | G | A | GTCTGGTTTT |
| **variant-base** | T | T | G | T | T | A | T | C | A | T | G |
| **var-length** | 1 | 1 | 1 | 1 | 1 | 1 | 1 | 1 | 1 | 1 | 10 |
| **one-based-reference-position** | 241 | 913 | 2500 | 3037 | 3267 | 5388 | 5986 | 6954 | 10193 | 10195 | 11287 |
| **iupac-variant-bases** | T | T | G | T | T | A | T | C | A | T | G |

| | 11 | 12 | 13 | 14 | 15 | 16 | 17 | 18 | 19 | 20 | 21 |
|---|---|---|---|---|---|---|---|---|---|---|---|
| **sampleID** | barcode73/ARTIC/medaka | barcode73/ARTIC/medaka | barcode73/ARTIC/medaka | barcode73/ARTIC/medaka | barcode73/ARTIC/medaka | barcode73/ARTIC/medaka | barcode73/ARTIC/medaka | barcode73/ARTIC/medaka | barcode73/ARTIC/medaka | barcode73/ARTIC/medaka | barcode73[illegible]ARTIC/medaka |
| **type** | snp | snp | snp | snp | snp | snp | del | del | snp | snp | snp |
| **reference-base** | C | C | T | A | C | G | ATACATG | TTTA | G | A | C |
| **variant-base** | T | T | C | G | T | T | A | T | A | T | A |
| **var-length** | 1 | 1 | 1 | 1 | 1 | 1 | 7 | 4 | 1 | 1 | 1 |
| **one-based-reference-position** | 14408 | 15279 | 16176 | 17236 | 19032 | 19525 | 21764 | 21990 | 23012 | 23063 | 23271 |
| **iupac-variant-bases** | T | T | C | G | T | T | A | T | A | T | A |

Sample_ID: barcode73/ARTIC/medaka
PANGO:None

Variant Status: confirmed

PHE-Label: E484K
WHO Label:
Alternate Names: E484K,

*Description:*

E484K containing sequences

*Information Sources* Source_1

Variant Calls Detected:

| Position | gene | protein | ref | variant | sample call | type | status |
|---|---|---|---|---|---|---|---|
| 23012 | S | surface glycoprotein | G | A | A | SNP | detect |

One based position - notes on sites detected in this sample.

The observed calls were:

- mutation_ref_calls: 0
- indel_ref_calls: 0
- mutation_calls: 1
- mutation_mixed_calls: 0
- indel_calls: 0
- no_calls: 0
- no_call_deletion: 0

Classification rules:

confirmed

mutations_required: 1

indels_required: 0

allowed_wildtype: 0

low_qc

mutations_required: 0

indels_required: 0

allowed_wildtype: 0

*Acknowledgements:*
*Curators:*Richard Myers

The complete list of variants of concern scanned included:
VUI-21FEB-04 VOC-20DEC-02 VUI-21FEB-01 VUI-21MAR-01 VUI-21MAY-01 VOC-20DEC-01 VOC-21APR-02 VUI-21FEB-03 VUI-21APR-01 E484K VUI-21JUL-01 VUI-21JAN-01 VUI-21MAR-02 VUI-21MAY-02 VUI-21JUN-01 VOC-21JAN-02 VUI-21APR-03 VOC-21FEB-02
