## Supplementary File 1 for "Real-time monitoring and analysis of SARS-CoV-2 nanopore sequencing with minoTour": 103_barcode86_artic_report.pdf

### Report for artic output of barcode86 generated 8th September 2021 14:18

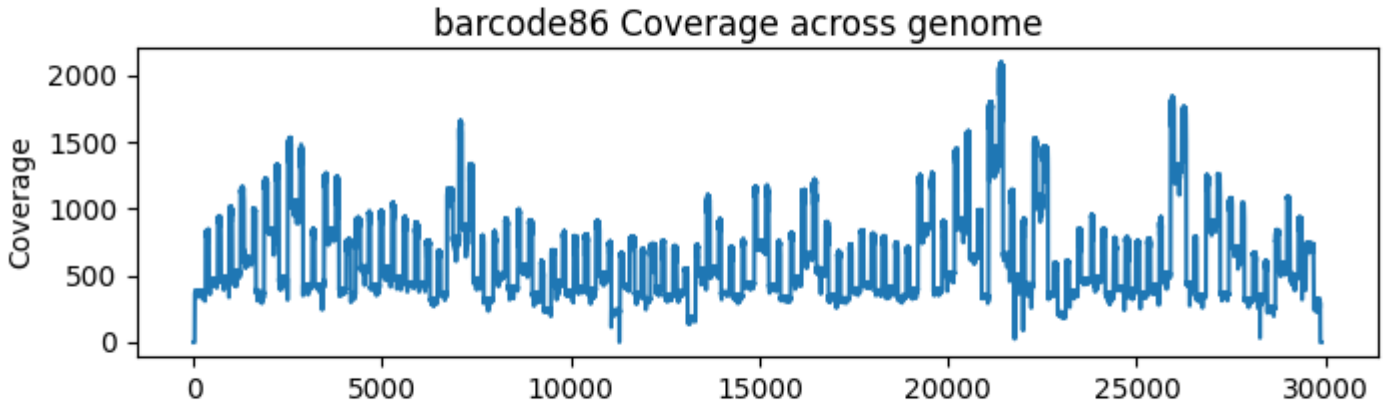

#### Artic variant summaries for barcode86

##### Variant of Concern Report for barcode86/ARTIC/medaka

If a genome is analysed, this will provide a report for the variant or variants found based on current PHE Variant Definitions. This analysis will report all possible VoCs and caution should be taken in interpretation of low coverage/quality genomes.

Reported VuI/VoCs:

| sample_id | phe-label | unique-id | status | mutation-ref-calls | mutation-mixed-calls | mutation-calls | indel-ref-calls | indel-calls | no-calls | no-calls-deletion |
|---|---|---|---|---|---|---|---|---|---|---|
| barcode86/ARTIC/medaka | VOC-20DEC-01 | denture-daughter | confirmed | 0 | 0 | 13 | 0 | 2 | 0 | 0 |

Observed mutations:

| | 0 | 1 | 2 | 3 | 4 | 5 | 6 | 7 | 8 | 9 | 10 |
|---|---|---|---|---|---|---|---|---|---|---|---|
| **sampleID** | barcode86/ARTIC/medaka | barcode86/ARTIC/medaka | barcode86/ARTIC/medaka | barcode86/ARTIC/medaka | barcode86/ARTIC/medaka | barcode86/ARTIC/medaka | barcode86/ARTIC/medaka | barcode86/ARTIC/medaka | barcode86/ARTIC/medaka | barcode86/ARTIC/medaka | barcode86/ARTIC/medaka |
| **type** | snp | snp | snp | snp | snp | snp | snp | snp | snp | del | ins |
| **reference-base** | C | C | C | T | C | C | C | C | T | GTCTGGTTTT | T |
| **variant-base** | T | T | T | C | T | T | A | T | C | G | TG |
| **var-length** | 1 | 1 | 1 | 1 | 1 | 1 | 1 | 1 | 1 | 10 | 2 |
| **one-based-reference-position** | 241 | 913 | 3037 | 3095 | 3177 | 3267 | 5388 | 5986 | 6954 | 11287 | 12803 |
| **iupac-variant-bases** | T | T | T | C | T | T | A | T | C | G | TG |

| | 11 | 12 | 13 | 14 | 15 | 16 | 17 | 18 | 19 | 20 | 21 |
|---|---|---|---|---|---|---|---|---|---|---|---|
| **sampleID** | barcode86/ARTIC/medaka | barcode86/ARTIC/medaka | barcode86/ARTIC/medaka | barcode86/ARTIC/medaka | barcode86/ARTIC/medaka | barcode86/ARTIC/medaka | barcode86/ARTIC/medaka | barcode86/ARTIC/medaka | barcode86/ARTIC/medaka | barcode86/ARTIC/medaka | barcode8[illegible] ARTIC/medaka |
| **type** | snp | snp | snp | snp | del | del | snp | snp | snp | snp | snp |
| **reference-base** | C | C | C | T | ATACATG | TTTA | A | C | A | C | C |
| **variant-base** | T | T | T | C | A | T | T | A | G | A | T |
| **var-length** | 1 | 1 | 1 | 1 | 7 | 4 | 1 | 1 | 1 | 1 | 1 |
| **one-based-reference-position** | 14408 | 15279 | 15960 | 16176 | 21764 | 21990 | 23063 | 23271 | 23403 | 23604 | 23709 |
| **iupac-variant-bases** | T | T | T | C | A | T | T | A | G | A | T |
