## Supplementary Information for "Real-time monitoring and analysis of SARS-CoV-2 nanopore sequencing with minoTour"

#### Supplementary figures

**A**

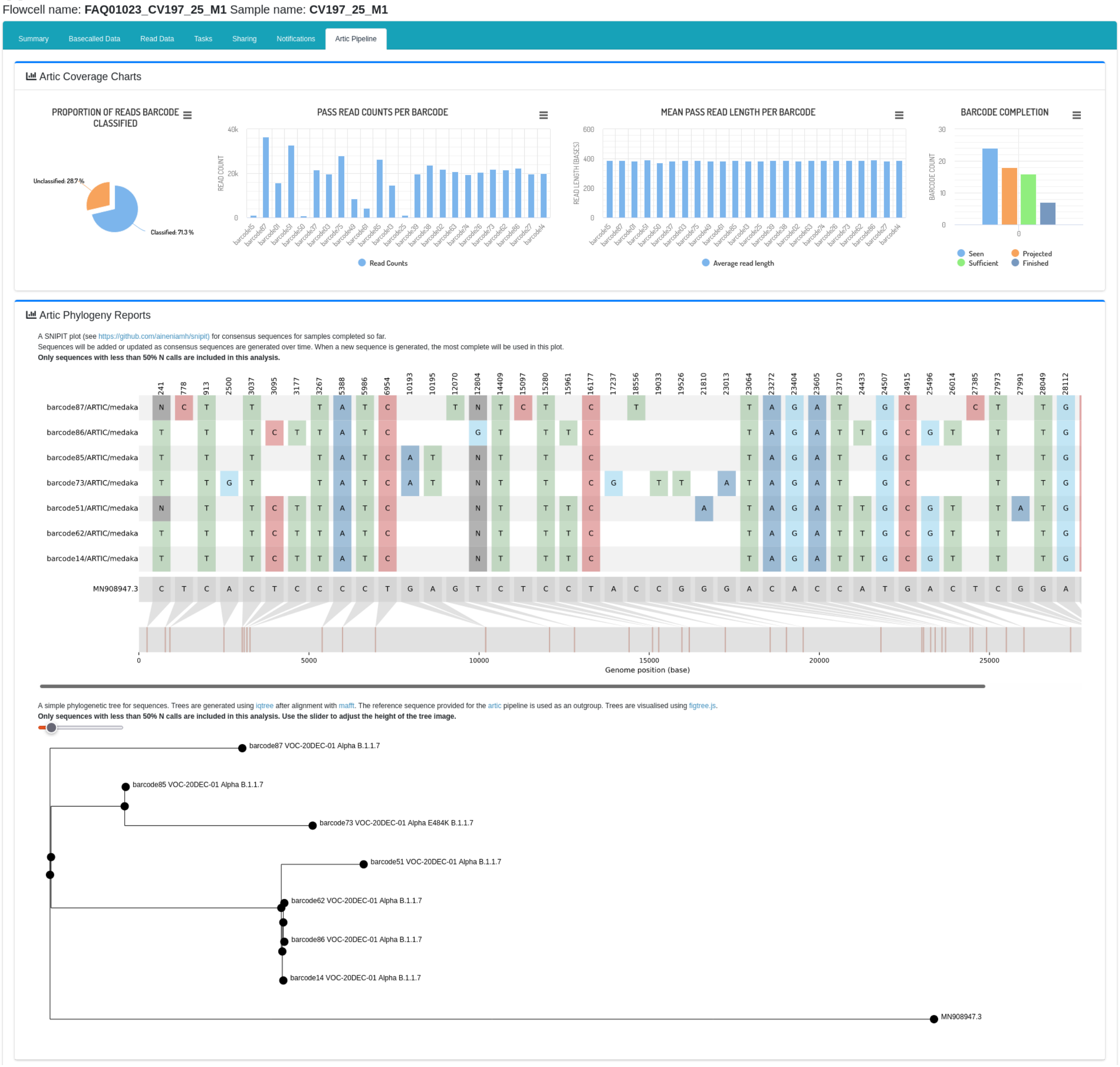

**B**

##### Pipeline firing conditions

20 X Coverage at 90 % of amplicons Submit choices

Show 10 entries Search:

| % of amplicons | X coverage | Delete |
|---|---|---|
| 90 | 20 | ✕ Delete |
| % of amplicons | X coverage | Delete |

Showing 1 to 1 of 1 entries

Previous 1 Next

##### Summary Table (Pass reads only)

Copy CSV PDF Show 10 rows Search: barcode

| Barcode | Sequence Name | Sequence Length | Read Count | Yield | Average Read Length | Coverage | # Success Amplicons | # Partial Amplicons | # Failed Amplicons | Mean of amplicons | Var of amplicons | Amplicon Std. Dev | Sufficient Coverage | Projected to Finish | Lineage | VoC Found |
|---|---|---|---|---|---|---|---|---|---|---|---|---|---|---|---|---|
| barcode15 | MN908947.3 | 29,903 | 1,137 | 440.68 kb | 388 | 14.45 | 16 | 4 | 79 | 13.4 | 1102.03 | 33.2 | false | false | Currently unknown | Not Tested |
| barcode87 | MN908947.3 | 29,903 | 36,440 | 14.12 Mb | 388 | 463.22 | 99 | 0 | 0 | 445.87 | 29669.97 | 172.25 | true | true | B.1.1.7 | VOC-20DEC-01 (confirmed) |
| barcode01 | MN908947.3 | 29,903 | 15,766 | 6.09 Mb | 386 | 199.52 | 71 | 9 | 19 | 188.82 | 60892.39 | 246.76 | false | false | Currently unknown | Not Tested |
| barcode51 | MN908947.3 | 29,903 | 32,764 | 12.81 Mb | 391 | 420.15 | 99 | 0 | 0 | 404.61 | 26610.79 | 163.13 | true | true | B.1.1.7 | VOC-20DEC-01 (confirmed) |
| barcode50 | MN908947.3 | 29,903 | 705 | 262.70 kb | 373 | 8.61 | 10 | 9 | 80 | 8.46 | 887.69 | 29.79 | false | false | Currently unknown | Not Tested |
| barcode37 | MN908947.3 | 29,903 | 21,563 | 8.31 Mb | 386 | 272.39 | 89 | 6 | 4 | 261.48 | 46975.64 | 216.74 | false | true | Currently unknown | Not Tested |
| barcode03 | MN908947.3 | 29,903 | 19,712 | 7.63 Mb | 387 | 249.89 | 92 | 7 | 0 | 243.16 | 24450.24 | 156.37 | true | true | Currently unknown | Not Tested |
| barcode75 | MN908947.3 | 29,903 | 28,058 | 10.87 Mb | 388 | 356.25 | 95 | 4 | 0 | 342.77 | 39780.85 | 199.45 | true | true | Currently unknown | Not Tested |
| barcode49 | MN908947.3 | 29,903 | 8,602 | 3.31 Mb | 384 | 108.39 | 56 | 13 | 30 | 108.21 | 22371.43 | 149.57 | false | false | Currently unknown | Not Tested |
| barcode61 | MN908947.3 | 29,903 | 4,319 | 1.67 Mb | 386 | 54.75 | 32 | 6 | 61 | 49.38 | 8000.96 | 89.45 | false | false | Currently unknown | Not Tested |
| Barcode | Sequence Name | Sequence Length | Read Count | Yield | Average Read Length | Coverage | # Success Amplicons | # Partial Amplicons | # Failed Amplicons | Mean of amplicons | Var of amplicons | Amplicon Std. Dev | Sufficient Coverage | Projected to Finish | Lineage | VoC Found |

Showing 1 to 10 of 24 entries (filtered from 25 total entries)

Previous 1 2 3 Next

Download all completed | Run all Incomplete | Re-Run All | Export report

**C**

#### Lineage Information for barcode87/ARTIC/medaka - B.1.1.7

Links to information on lineage B.1.1.7

PANGO Info | Outbreak Info

#### Variant of Concern Report for barcode87/ARTIC/medaka

If a genome is analysed, this will provide a report for the variant or variants found based on current PHE Variant Definitions. This analysis will report all possible VoCs and caution should be taken in interpretation of low coverage/quality genomes.

Reported VuI/VoCs:

| sample_id | phe-label | unique-id | status | mutation-ref-calls | mutation-mixed-calls | mutation-calls | indel-ref-calls | indel-calls | no-calls | no-calls-deletion |
|---|---|---|---|---|---|---|---|---|---|---|
| barcode87/ARTIC/medaka | VOC-20DEC-01 | denture-daughter | confirmed | 0 | 0 | 13 | 0 | 2 | 0 | 0 |

Observed mutations:

| type | snp | snp | snp | snp | snp | snp | snp | del | snp | snp | snp | snp | snp | snp | del |
|---|---|---|---|---|---|---|---|---|---|---|---|---|---|---|---|
| **reference-base** | T | C | C | C | C | C | T | GTCTGGTTTT | G | C | T | C | T | C | ATACATG |
| **variant-base** | C | T | T | T | A | T | C | G | T | T | C | T | C | T | A |
| **var-length** | 1 | 1 | 1 | 1 | 1 | 1 | 1 | 10 | 1 | 1 | 1 | 1 | 1 | 1 | 7 |
| **one-based-reference-position** | 778 | 913 | 3037 | 3267 | 5388 | 5986 | 6954 | 11287 | 12070 | 14408 | 15096 | 15279 | 16176 | 18555 | 21764 |
| **iupac-variant-bases** | C | T | T | T | A | T | C | G | T | T | C | T | C | T | A |

| type | del | snp | snp | snp | snp | snp | snp | snp | snp | snp | snp | snp | del | mnp | mnp-snp | mnp-snp |
|---|---|---|---|---|---|---|---|---|---|---|---|---|---|---|---|---|
| **reference-base** | TTTA | A | C | A | C | C | T | G | T | C | G | A | TA | GAT | A | T |
| **variant-base** | T | T | A | G | A | T | G | C | C | T | T | G | T | CTA | T | A |
| **var-length** | 4 | 1 | 1 | 1 | 1 | 1 | 1 | 1 | 1 | 1 | 1 | 1 | 2 | 3 | 1 | 1 |
| **one-based-reference-position** | 21990 | 23063 | 23271 | 23403 | 23604 | 23709 | 24506 | 24914 | 27384 | 27972 | 28048 | 28111 | 28270 | 28280 | 28281 | 28282 |
| **iupac-variant-bases** | T | T | A | G | A | T | G | C | C | T | T | G | T | CTA | T | A |

D

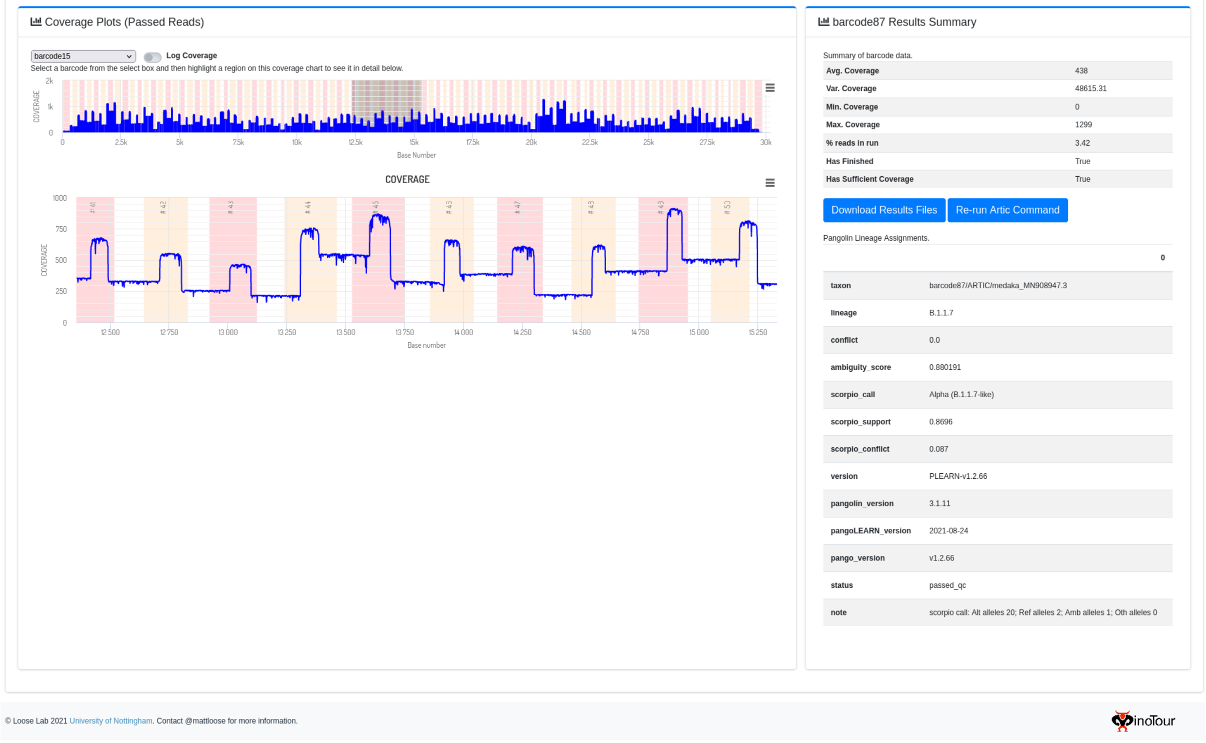


**Supplementary figure 1 -** Screenshot of the Artic tab in minoTour. **A)** The top section of the tab deals with visualising overall run metrics, such as the proportion of mapped reads classified as a barcode in a run, the read count per barcode, the mean read length and the total number of amplicons that have over 20X coverage, over 1X coverage and 0X coverage. All generated consensuses have a SNIPIT report generated for them, and a phylogenetic tree generated. **B)** The next section of the tab displays the conditions required for a sample to have the ARTIC medaka pipeline run on it's accumulated reads. The default for a sample to be analysed is displayed, with 90% of amplicons on the sample at 20X. A sortable and searchable summary table displays information about each sample currently identified in the run. Samples that have been run are colour coded as green rows. **C)** This section deals with information about a specific sample chosen from the table in B) by clicking on it. If the sample has sufficient data and the medaka pipeline has been run on it, a report generated by the aln2type tool is displayed, confirming Lineage and SNPs seen in the consensus. **D).** The coverage is displayed for the genome selected from the table in B), along with the ARTIC lineage report.

### Supplementary files.

#### Supplementary file 1

PDF reports generated by the ARTIC tab in minoTour, bundled and compressed into a tar.gz file. Each barcode has a report generated, showing a simple line chart of coverage along the genome, and if the barcode has been run through the ARTIC medaka pipeline, the aln2type output report is included. A PDF summary for the whole run is also included, containing the mean read length and count per barcode, the SNIPIT plot of the run and the phylogenetic tree.

#### Supplementary file 2

CSV of lineage assigned to each sample at the FR, RU and SA timepoints for both the medaka and nanopolish generated consensus sequences. Cells marked N/A did not have sequences for that time point for both medaka and nanopolish, and cells marked None did not generate a lineage after running pangolin on them.

#### Supplementary file 3

Output CSV of nextclade (https://clades.nextstrain.org) web app on all our medaka consensus sequences.

#### Supplementary file 4

Output CSV of nextclade (https://clades.nextstrain.org) web app on all our nanopolish consensus sequences.
